# Identification of phenotype-specific networks from paired gene expression-cell shape imaging data

**DOI:** 10.1101/2021.02.11.430597

**Authors:** Charlie George Barker, Eirini Petsalaki, Girolamo Giudice, Julia Sero, Emmanuel Nsa Ekpenyong, Chris Bakal, Evangelia Petsalaki

**Affiliations:** European Molecular Biology Laboratory-European Bioinformatics Institute, Hinxton CB10 1SD, UK; University of Bath, Claverton Down, Bath BA2 7AY, UK; Institute of Cancer Research, 237 Fulham Road, London, SW3 6JB, UK

## Abstract

The morphology of breast cancer cells is often used as an indicator of tumour severity and prognosis. Additionally, morphology can be used to identify more fine-grained, molecular developments within a cancer cell, such as transcriptomic changes and signaling pathway activity. Delineating the interface between morphology and signaling is important to understand the mechanical cues that a cell processes in order to undergo epithelial-to-mesenchymal transition and consequently metastasize. However, the exact regulatory systems that define these changes remain poorly characterised. In this study, we employ a network-systems approach to integrate imaging data and RNA-seq expression data. Our workflow allows the discovery of unbiased and context-specific gene expression signatures and cell signaling sub-networks relevant to the regulation of cell shape, rather than focusing on the identification of previously known, but not always representative, pathways. By constructing a cell-shape signaling network from shape-correlated gene expression modules and their upstream regulators, we found central roles for developmental pathways such as WNT and Notch as well as evidence for the fine control of NFkB signaling by numerous kinase and transcriptional regulators. Further analysis of our network implicates a gene expression module enriched in the RAP1 signaling pathway as a mediator between the sensing of mechanical stimuli and regulation of NFkB activity, with specific relevance to cell shape in breast cancer.

## Introduction

The study of cancer has long been associated with changes in cell shape as morphology can be a reliable way to sub-type cancer and predict patient prognosis [1]. Recent research has implicated cellular morphology in more than just a prognostic role in cancer, with shape affecting tumour progression through the modulation of migration, invasion and overall tissue structure [2][3]. The unique mechanical properties of the tumour tissue (primarily driven by changes in cell shape and the extracellular matrix) are hypothesised to contribute to the ‘stem cell niche’ of cancer cells that enables them to self-renew as they do in embryonic development [4]. Cell morphology and tumour organisation have been found to be a factor in modulating the intra-cellular signaling state through pathways able to integrate mechanical stimuli from the extracellular environment [5][6][7][8]. The discovery of mechanosensitive pathways in various tissues has revealed a complex interplay between cell morphology and signaling [9]. Further studies have revealed that cell morphology can also be a predictor of tumorigenic and metastatic potential as certain nuclear and cytoplasmic features enhance cell motility and spread to secondary sites [1], aided by the Epithelial to Mesenchymal Transition (EMT). This process is the conversion of epithelial cells to a mesenchymal phenotype, which contributes to metastasis in cancer and worse prognosis in patients [10].

Breast cancer is the most common cancer among women, and in most cases treatable with a survival rate of 99% among patients with a locally contained tumour. However, among those patients presenting with a metastatic tumour this rate drops to 27% [11]. During the development of breast cancer tumours, cells undergo progressive transcriptional and morphological changes that can ultimately lead towards EMT and subsequent metastasis [12][13][1]. Breast cancer sub-types of distinct shapes show differing capacities to undergo this transition. For example, long and protrusive basal breast cancer cell lines are more susceptible to EMT [14] with fewer cell-to-cell contacts [15]. Luminal tumour subtypes on the other hand, are associated with good to intermediate outcomes for patients [15] and have a clear epithelial (or ‘cobblestone’) morphology with increased cell-cell contacts [16]. It is evident that cell morphology plays significant roles in breast cancer and a deeper understanding of the underlying mechanisms may offer possibilities for employing these morphology determinant pathways as potential therapeutic targets and predictors of prognosis.

Signaling and transcriptomic programs are known to be modulated by external physical cues in the contexts of embryonic development [17], stem-cell maintenance [18][19] and angiogenesis [20]. Numerous studies have flagged NFkB as a focal point for mechano-transductive pathways in various contents [21][22][23][24], but gaps in our knowledge remain as to how these pathways may interact and affect breast cancer development. Sero and colleagues studied the link between cell shape in breast cancer and NFkB activation by combining high-throughput image analysis of breast cancer cell lines with network modelling [25]. They found a relationship between cell shape, mechanical stimuli and cellular responses to NFkB and hypothesised that this generated a negative feedback loop, where a mesenchymal-related morphology enables a cell to become more susceptible to EMT, thus reinforcing their metastatic fate. This analysis was extended by [26], who combined cell shape features collected from image analysis with microarray expression data for breast cancer cell lines to create a shape-gene interaction network that better delineated the nature of NFkB regulation by cell shape in breast cancer. This approach was limited as it only correlates single genes with cell shapes, thus relying on the assumption that a gene’s expression is always a useful indicator of its activity [27]. Furthermore, the authors rely on a list of pre-selected transcription factors of interest and as such the approach is not completely data-driven and hypothesis free. Given our knowledge of the multitude of complex interacting signaling pathways in development and other contexts, it is safe to assume that there are many more players in the regulation of cancer cell morphology that have yet to be delineated [28][29][30][31]. Furthermore, how exactly extracellular mechanical cues are ‘sensed’ by the cell and passed on to NFkB in breast cancer is not clearly understood. From this it is clear that an unbiased approach is needed to identify novel roles for proteins in the interaction between cell shape and signaling.

Here we identify a data-derived cellular signaling network, specific to the regulation of cell shape beyond NFkB, by considering functional co-expression modules and cell signaling processes rather than individual genes. To this end, we have developed a powerful network-based approach to bridge the gap between widely available and cheap expression data, signaling events and large-scale biological phenotypes such as cell shape (Figure 1A). By organizing expression data into context-specific modules we leverage the transcriptome’s propensity to be regulated in regulons, thereby aiding the inference of signaling activity. This, combined with the use of feature-correlated modules, allows for the analysis of complex phenotypes, defined by multiple features. The resulting signaling network was validated using data from cell treatments with kinase inhibitors [32] and revealed further regulation of NFkB’s utility in modulating mechanotransduction in response to morphological changes in breast cancer.

**Figure 1:**
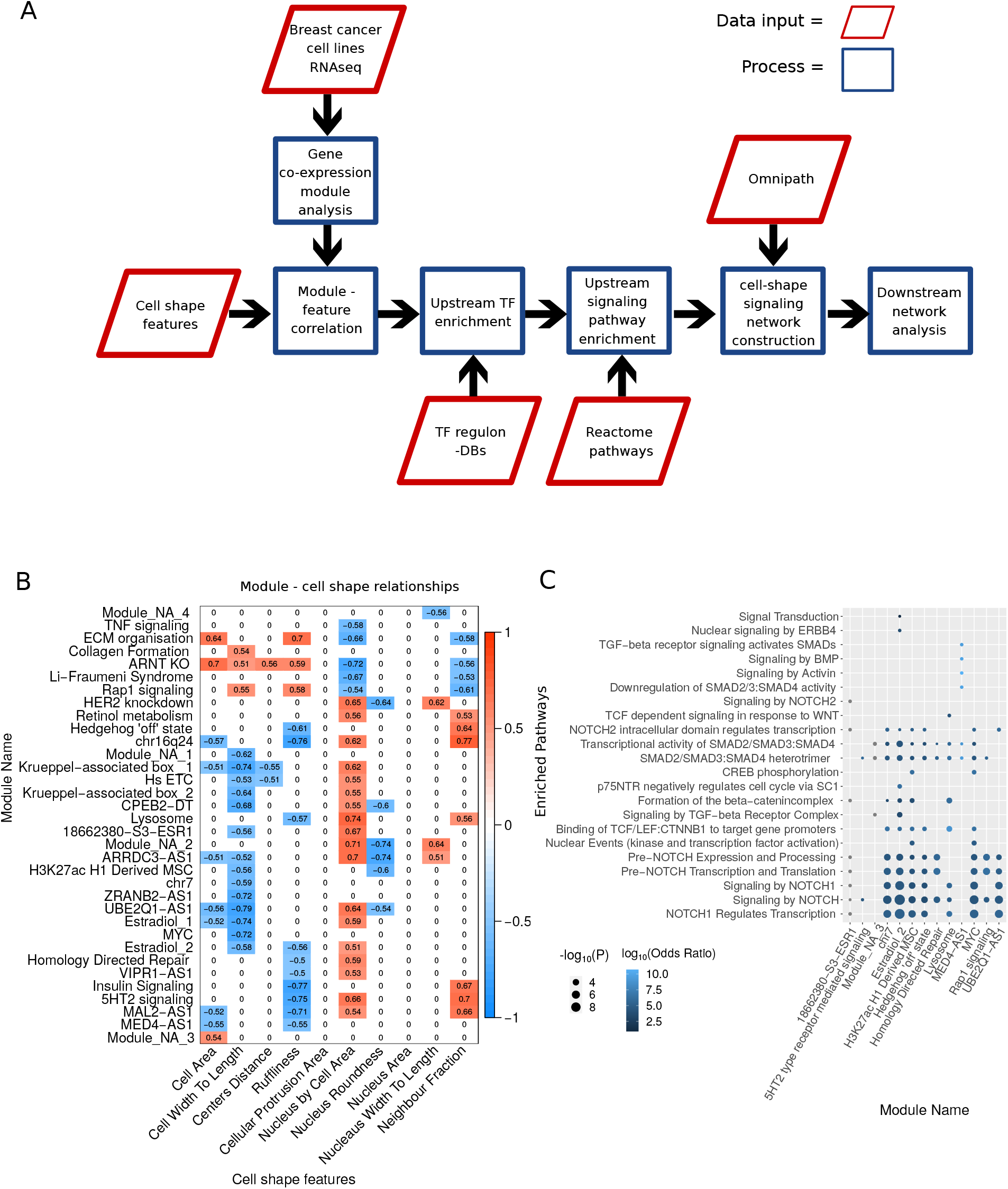
**A**. Schematic illustrating the steps involved in phenotype-specific network construction. Gene expression modules are identified by integrating cell shape variables (derived from imaging data) with RNAseq data from breast cancer cell lines. These gene expression modules are correlated with specific cell shape features to find morphologically relevant modules. Next, transcription factors (TFs) are identified whose targets significantly overlap with the contents of the expression modules. These TFs are used to identify pathways regulating the gene expression modules, which are then integrated to form a contiguous network using PCSF. **B**. Heatmap of significantly correlated gene expression module eigengenes with cell shape features. Non-significant interactions were set to 0 for clarity. **C**. Dot plot illustrating the enrichment of pathways among TFs found to regulate gene expression modules. The x axis shows the module names (as defined by Supplementary Table 3) and the y axis shows the signaling pathways found to be significantly (P<0.01) enriched in the TFs that regulate the given module (as defined by Supplementary Table 5). The y axis is arranged such that the terms with the highest combined odds ratio are at the bottom. Size of the dot represents the -log10(P) and the colour indicates a log 10 transformation of the odds ratio.

## Results

### Identification of gene co-expression modules correlated with cell shape features

We first sought to identify gene expression modules (GEMs) that are relevant to the regulation of cell shape. To this end, we used Weighted Gene Correlation Network Analysis (WGCNA) [33] on bulk RNA-seq expression data from 13 breast cancer and one non-tumorigenic epithelial breast cell lines to identify gene co-expression modules correlated with 10 specific cell shape variables [26] (Methods). These described the size, perimeter and texture of the cell and the nucleus (n = 75,653). Of 102 GEMs (Figure S1A), 34 were significantly correlated (P<0.05; Student’s T-test, Pearson Correlation; Supplementary Table 1) with one of 8 cell shape features (Figure 1B). A full list of the genes within the identified modules are summarized in Supplementary Table 2.

We used EnrichR and their suite of gene set libraries [34] to functionally annotate and label some of the modules using enrichment of genes contained within them. We found that the ‘RAP1 signaling’ module is also enriched for terms such as VEGF signaling and hemostasis, while the ‘Insulin Signaling’ module is also enriched for cell-cell communication and the ‘ECM organisation’ module is also enriched in terms such as axon guidance and EPH-Ephrin signaling (Supplementary Table 3). Modules that are most correlated with all features are the ‘ARNT KO’ module, ‘ARRDC3AS1’ module and the ‘ECM organisation’ module (see Figure S1B). Modules that could not be annotated with informative terms were designated ‘module non-annotated (NA) 1, 2, 3 etc.

### Transcription factor analysis of cell shape gene co-expression modules reveals the signaling pathways that regulate them

To link these expression modules to the intra-cellular signaling network, we considered both the regulation of modules as transcriptional units as well as the signaling pathways that significantly regulate the identified regulons. Specifically, we first found 17 transcription factor (TF) regulons, as defined in the database TRRUST v2 [35], to be significantly enriched (P<0.1; Fisher’s exact test) in our modules (Supplementary Table 4). We therefore consider these TFs as potentially relevant for the regulation of cell shape features and their activity levels as a read-out of cell signaling activity in these cells. These TFs include the EMT antagonist FOXA1 [36], and HOXB7 [37] and ZFP36 [38].

To extend this further, we sought to investigate the pathways responsible for regulating the identified TFs, and by extension the gene expression modules. For this analysis, we also include ENCODE and ChEA Consensus TFs from ChIP-X [39],DNA binding preferences from JASPAR[40][41], TF protein-protein interactions and TFs from ENCODE ChiP-Seq [42] to get a more comprehensive picture of the pathways involved in regulation of cell morphology. Using the identified TFs (Supplementary Table 5) we then used EnrichR [34] to perform a Reactome signaling pathway [43] enrichment analysis. Results from this analysis showed that many identified gene expression modules were regulated by common signaling pathways, with 6 modules sharing pathways associated with downstream signaling and regulation of NOTCH (Figure 1C). In order to ensure that our approach is not biased to any particular pathway, we repeated our approach on 1,000 resampled GEMs, and created pathway-specific null distributions for each identified pathway. All pathways we identified from morphology-correlated modules had significantly lower p-values than randomised modules (FDR adjusted P<0.05). The only exceptions were one association with “Signaling by NOTCH” and modules associated with “Signal Transduction”, a spurious pathway containing the complete intra-cellular signaling system (Supplementary Table 6).

### Clustering based on morphology reveals distinctive cell-line shapes

To understand key differences in expression patterns and gene regulation between morphologically distinct breast cancer cell lines, we clustered them based on 10 morphological features including area, ruffliness, protrusion area and neighbour frequency and performed differential expression analysis between the identified clusters (Figure 2A). Cluster A is more heterogeneous in its morphology, containing the non-tumorigenic mammary epithelial cell line MCF-10A as well as cell lines from both luminal and basal breast cancer subtypes. Clusters B and C are more distinctly shaped, roughly composed of luminal and basal cell lines respectively with the exception of HCC1954, which was clustered morphologically with luminal subtypes while being characterised as basal. The basal-like cluster is most morphologically distinct from cluster A, but also differs from the luminal-like cluster in that it has a lower nuclear/cytoplasmic area (0.133 ± 0.05 [mean ± SD]), higher ruffliness (0.235 ± 0.12) and lower neighbour fraction (0.258 ± 0.22). The luminal-like cluster had a higher nuclear/cytoplasmic area (0.186 ± 0.1; P<0.001), lower ruffliness (0.213 ± 0.14; P<0.001), and a higher neighbour fraction (0.338 ± 0.26; P<0.001, One-way ANOVA; Tukey HS, n = 75,653). The neighbour fraction feature corresponds to the fraction of the cell membrane that is in contact with neighbouring cells. The lower number of cell-cell contacts in basal-like breast cancer cell lines are indicative of more mesenchymal features associated with worse prognosis due to metastasis. Increased cell-cell contacts in both the luminal-like cluster and the more heterogeneous cluster A correspond to ‘cobblestone’ epithelial morphology. Interestingly, these groups are closely aligned with the expression of the cell adhesion protein, N-cadherin (Figure 2A), the expression for which is closely associated with a migratory and metastatic phenotype [44]. Representative images of the morphologically clustered cell lines are shown along with the clustering heatmap in Figure 2A (complete dataset of images provided online; https://datadryad.org/stash/dataset/doi:10.5061/dryad.tc5g4).

**Figure 2:**
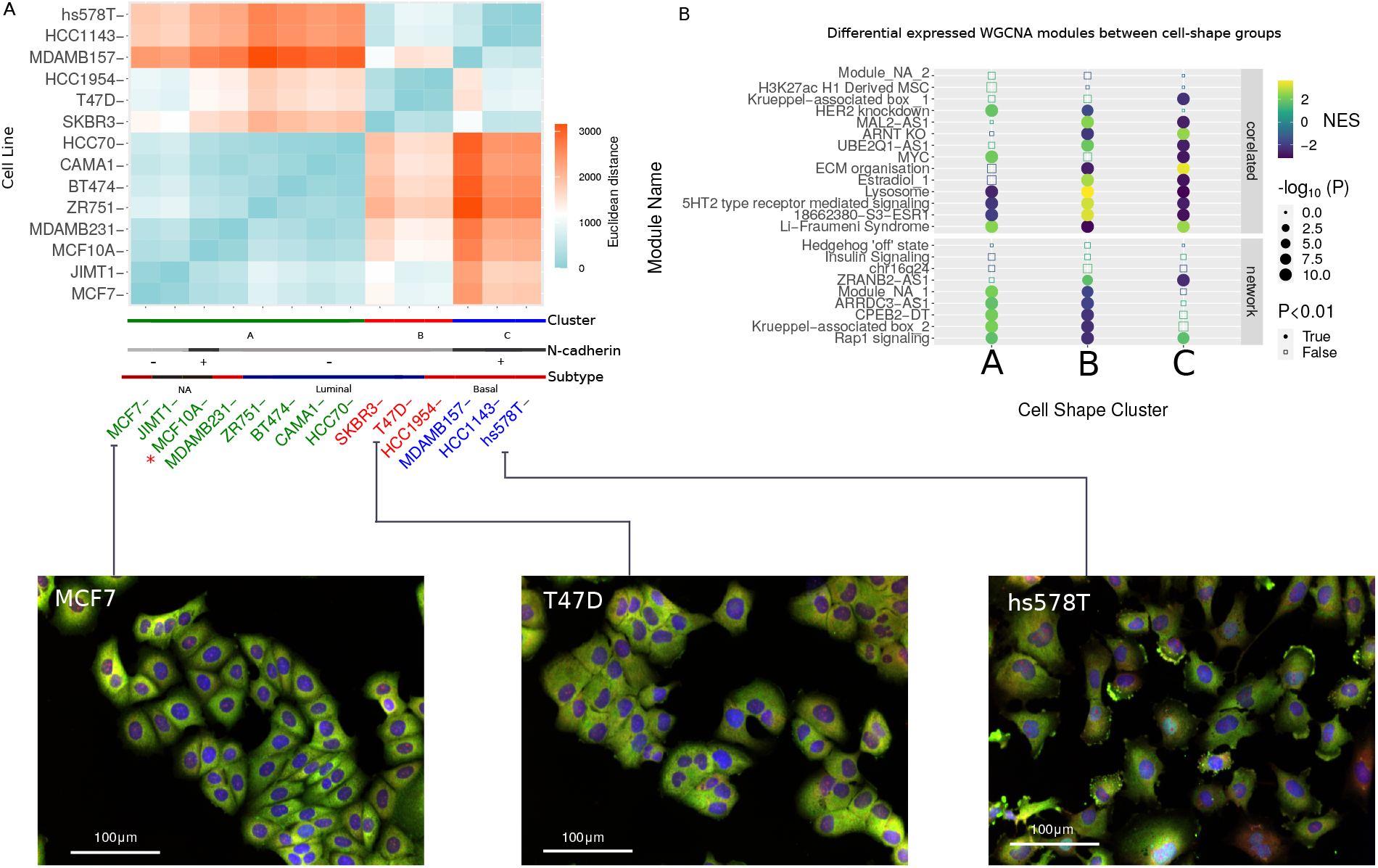
**A**. Heatmap of Euclidean distance between cell lines for shape features to illustrate clusters arising from k-means method. The coloured lines on the bottom show the assigned cluster and the cadherin expression and assigned canonical cancer subtype. **B**. Dotplot showing the enrichment of gene expression modules in the different cell line clusters. Along the y axis are the names of the clusters, faceted by whether they are included in the PCSF derived regulatory network on the bottom and whether they are correlated with cell shape variables, but not included in the network on the top. The x axis shows the cell shape clusters, with letters corresponding to the groups in Figure 2A (A - a heterogeneous mix of breast cancer subtypes, B - luminal-like cell lines and C - basal-like cell lines.). Dots are coloured based on the normalised enrichment value, with down-regulated modules in blue, and the up-regulated modules in yellow. Size corresponds to significance (-log10(P)) with the shape illustrating which changes are significant (adjusted P<0.01, Benjamini-Hochberg). **C**. Images (see Methods) showing morphology of representative cell lines from each respective cluster.Colours indicate labeling with DAPI (blue), Alexa 488 (green) and DHE (red).

Using the identified groups of cell lines in the previous step, differential expression analysis and transcription factor activity analysis was used to study gene regulation signatures specific to cell line morphological clusters. The results are shown in Supplementary Table 7, with gene set enrichment analysis showing upregulation of genes involved in the extracellular matrix, collagens, integrins and angiogenesis in the basal-like cluster. Significantly enriched terms (P<0.05) in downregulated genes include ‘fatty acid and beta-oxidation’ and ‘ERBB network pathway’. In the genes upregulated in the luminal-like cluster, we observed enrichment of terms such as ‘hallmark-oxidative phosphorylation’. Downregulated genes were enriched in ‘integrin-1 pathway’, ‘core matrisome’ and genes linked to ‘hallmark epithelial-mesenchymal transition and migration’. For the remaining B/L group, the term with the highest normalized enrichment score was ‘targets of the transcription factor Myc’ followed by terms associated with ribosomal RNA processing. Down-regulated terms include ‘cadherin signaling pathway’ (Supplementary Table 7).

We also calculated the differential expression for the WGCNA gene expression modules and found distinct patterns of expression between luminal-like and basal-like clusters of cell lines (Figure 2B). Among these, the RAP1 Signaling module is upregulated in basal-like clusters and downregulated in luminal-like clusters. This is consistent with the fact that this gene expression module is negatively correlated with neighbour fraction, a feature that is observed to decrease in mesenchymal-like cell shapes [15]. Other modules whose expression distinguishes basal-like from luminal-like include the MAL2-AS1 module (enriched in desmosome assembly), ARNT/KO module (enriched in TNF-signaling by NFkB) and ECM organisation module (enriched in focal adhesion proteins - see Supplementary Table 3).

To link the observed gene expression differences to cell signaling we used the tool DOROTHEA [45] to calculate transcription factor activities, as their modulation is one of the main results of cell signaling processes. We corroborated that the heterogenous B/L group had significantly activated Myc levels. In the luminal-like cluster, ESRRA (estrogen-related receptor *α*) is the most significantly overrepresented regulome, followed by EHF, KLF5 and ZEB2. Under-represented regulomes include KLF4, SMAD4, SMAD2, SOX2 and RUNX2. For the basal-like cluster, the regulome with the highest normalised enrichment score is SOX2, as well as Musculin and HOXA9. Downregulated regulomes include ZEB2, Myc, ESRRA and KLF5 (Figure S2D).

### Assembly of a data-driven cell shape regulatory network

In order to integrate our data-driven GEMs with signaling pathways, we used the Prize Collecting Steiner Forest (PCSF) algorithm [46]. This is an approach that aims to maximise the collection of ‘prizes’ associated with inclusion of relevant nodes, while minimizing the costs associated with edge-weights in a network. This allowed for the integration of the WGCNA modules, the Reactome pathways that regulate them, the TRRUST transcription factors and the differentially expressed DOROTHEA regulons into a contiguous regulatory network describing the interplay between cell shape and breast cancer signaling. The network used for this process was extracted from the database OmniPath [47] to provide a map of the intracellular signaling network described as a signed and directed graph. We incorporated identified GEMs into the network by interlinking them as nodes with the relevant TFs and signalling pathways.

The resulting network of 691 nodes included 97.11% of the genes identified by our analysis (Figure 3A). The new proteins that were included by the PCSF algorithm to maximise prize collection showed gene set enrichment of common terms (Pathways from Panther [48]) relative to the original prizes (including WNT, EGF, Angiogenesis, Ras, Cadherin and TGF-*β* pathways), but also included are some new terms (VEGF, Integrin and Endothelin pathways) (P<0.001; Figure 3B).

**Figure 3:**
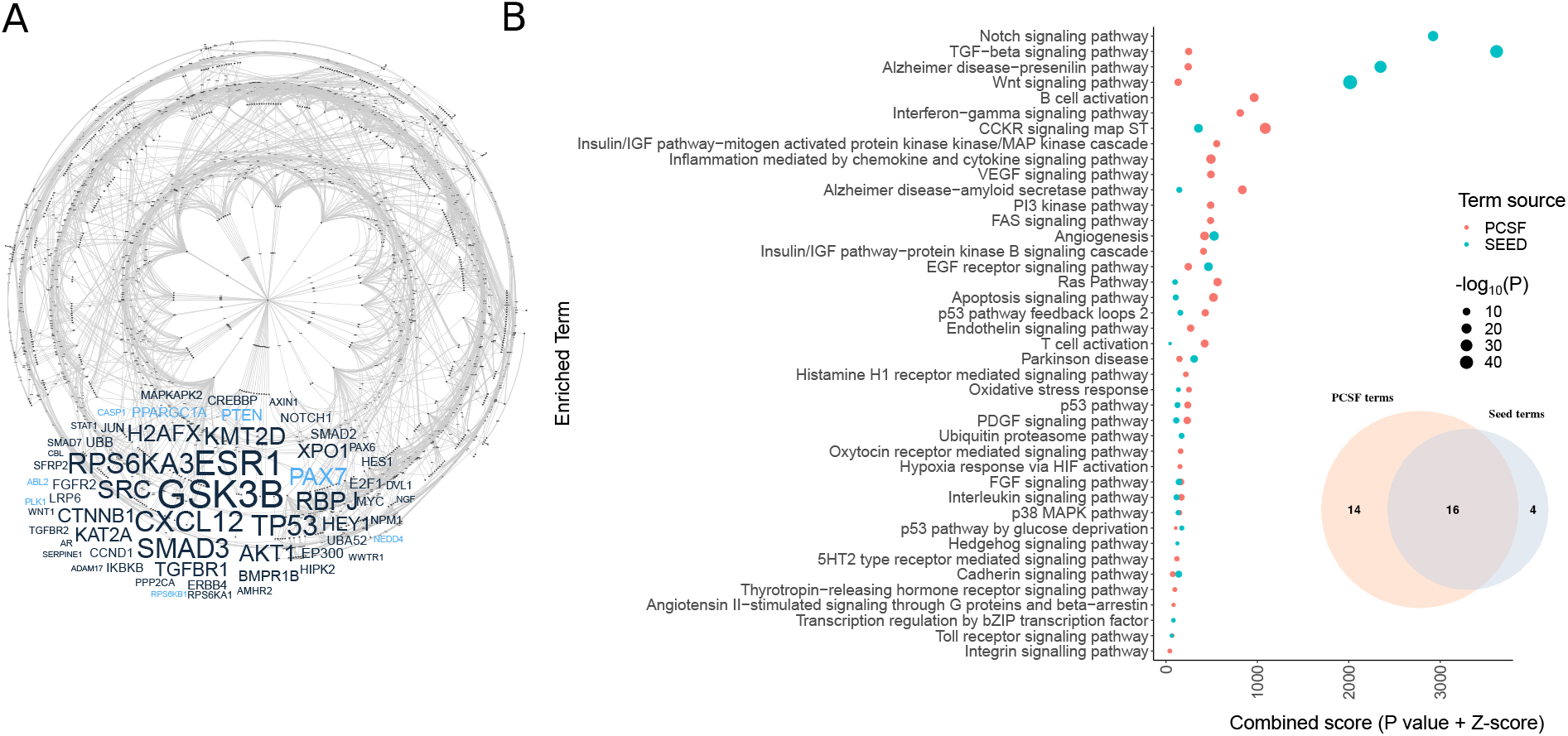
**A**. Network derived from integrating enriched pathways and transcription factor regulons within cell shape correlated gene expression modules. Below is a word cloud illustrating the top 50 nodes with highest betweenness centrality in the network, with those nodes belonging to the original seeds coloured black, and those genes included by the PCSF step coloured light blue. The size of the word in the word cloud corresponds to the nodes with the highest betweenness centrality. **B**. Gene set enrichment (Panther DB 2016) of original seed nodes and the nodes included by PCSF with the dot size indicating the level of significance (-Log10P) of the term enrichment. Blue nodes represent enrichments from proteins that where used as inputs for the PCSF algorithm and red nodes are those that were included because of it. Inset is a Venn diagram showing the overlap in enriched terms between the seed nodes and the terms included by PCSF.

Studying the network properties of our PCSF-derived regulatory network we find that the degree distribution is typical for a biological network (Figures S3A-C). The proteins in the network can be ranked by betweenness centrality to disseminate them based on network importance. Nodes with high centrality lie between many paths and can control information flow. Proteins with the highest centrality are primarily prizes (GSK3*b*, ESR1, p53, SMAD3 - Figure S3D) indicating that the PCSF solution was not achieved by the inclusion of new hub proteins that are not of interest to our analysis. That being said, a small minority of high centrality nodes were not in the original prizes, implicating them as mediating the cross talk between pathways identified in figure 1C. These include the proteins PAX7, PTEN and PPARGC1a.

### Small-molecule inhibitors targeting kinases in our network significantly perturb cell morphology

To validate our network, we used an independent dataset to evaluate whether perturbing the function of kinases within our predicted network would produce a significant effect on morphological features. For this, we used the Broad Institute’s Library of Integrated Network-based Cellular Signatures (LINCS) small molecule kinase inhibitor dataset [32]. Here, they measured morphological changes in the breast cancer cell line HS578T in response to various small molecule kinase inhibitors using high through-put imaging techniques [49]. The morphological variables measured in this data-set are mostly analogous to the ones used to construct the network, however there are some discrepancies which we used as negative controls to ensure our network was phenotype-specific.

We combined this with data from a target affinity assay [50] describing the binding affinities of small molecules to kinases. This enabled us to sort the kinase inhibitors into those that target proteins we predict regulate cell shape (through their inclusion in the PCSF derived network) and those that do not. Figure 4A illustrates that there is a statistically significant (n = 37, Wald test P<0.05) deviation from the control between drug treatments targeting kinases within the predicted network and those targeting other proteins for cytoplasmic area, cytoplasmic perimeter, nucleus area, nucleus length, nucleus width and nucleus perimeter. This difference is insignificant for features that were not correlated with gene expression modules in our initial analysis (such as number of small spots in the cytoplasm and nucleus, and nuclear compactness), indicating that our network is phenotype-specific to the features used in network generation. We also repeated this analysis in other cell lines (SKBR3, MCF7 and non-tumorigenic mammary cell line MCF10A) with results with limited statistical significance (Supplementary table 8). We additionally used a positive control where the control cells had been treated with TRAIL (TNF-related apoptosis-inducing ligand) in order to ensure that the observed morphological effects were not caused by apoptotic factors (S4B, Supplementary Table 8).

**Figure 4:**
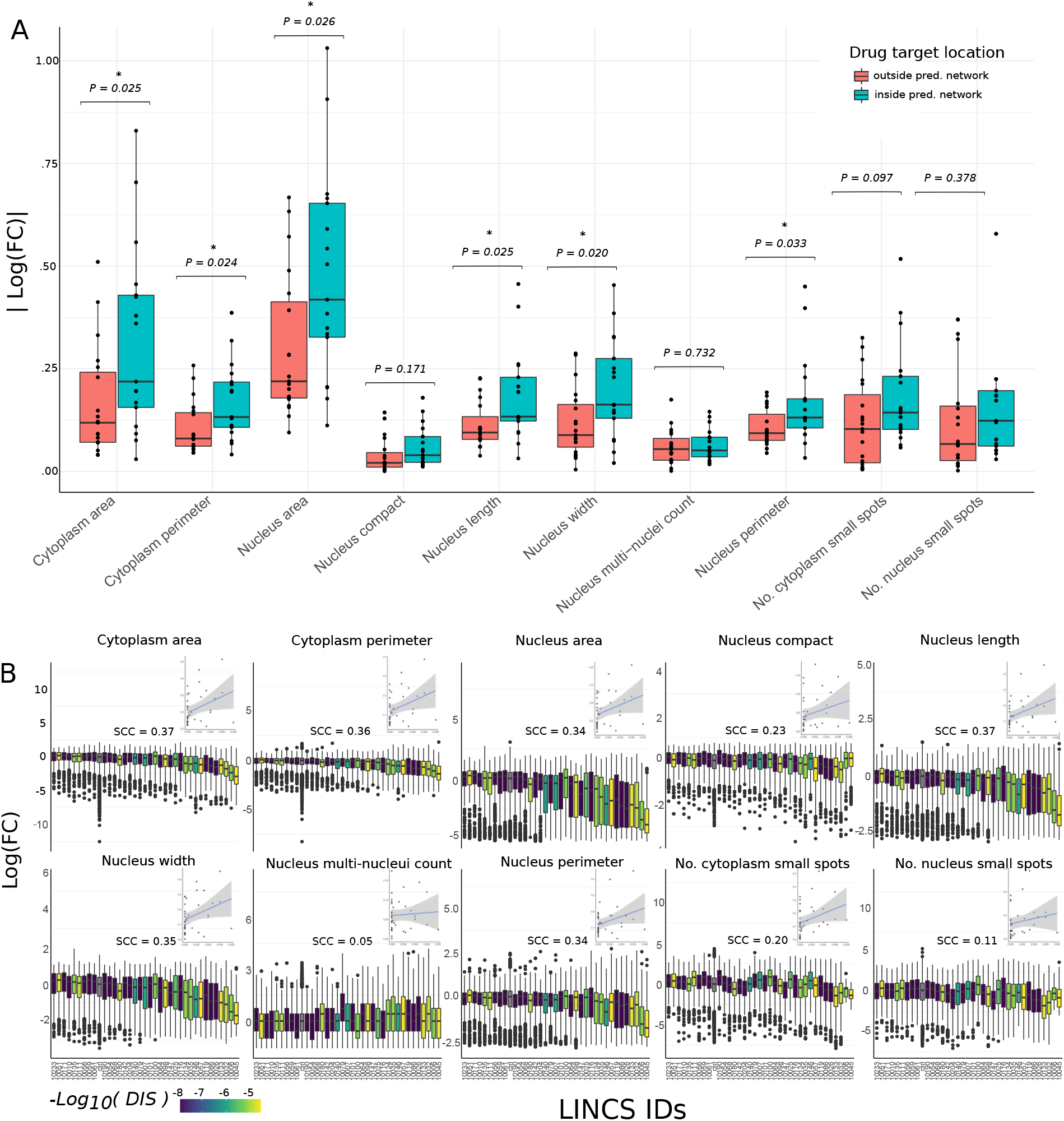
**A**. Box plots showing the absolute log(10) fold changes after treatment with a drug relative to a control for each cell shape variable. The drugs are grouped depending on whether they target kinases within the predicted regulatory network (blue) and those targeting other kinases not predicted to be associated with cell shape (red). P values (Welch Two Sample t-test) are showing with stars indicating significance. **B**. Bar plot showing the absolute difference in log fold changes of cell shape variables after treatment with a drug relative to a control. Here, each drug is shown separately (with the LINCs ID shown on the x axis) and coloured based on the drug influence score (DIS) and each data-point represents a single cell. Inset are plots showing the correlation between this influence score and the difference between mean treated cells and mean control cells in each of the 10 measured cell shape features for each drug. Spearman correlation coefficients are shown above the inset plots.

Interestingly, there is greater variance in the effect size for kinase inhibitors targeting proteins contained within the predicted regulatory network than those outside. The individual effect on cell morphology for each drug is shown in S4A-B. We hypothesised that it was the network properties of kinases within our network that dictated their effect on morphological features, with some targets being on the periphery of our predicted network and therefore having limited influence over the regulation of cell shape. To test this, we studied the extent to which the effect of a kinase inhibitor was correlated with the combined centrality of its targets as defined by our network. For this we used the centrality algorithm PageRank (Brin and Page, 1998) and accounted for off-target effects of the kinase inhibitors using the Szymkiewicz-Simpson index (describing the overlap of a kinase inhibitor’s targets and the proteins that constitute the network - Methods).

Figure 4B shows moderate correlations between target centrality and the effect size for each feature, illustrating that kinase inhibitors targeting proteins with high centrality in our network modulate cell shape more than inhibitors with peripheral targets. As with studying the effect of targeting kinases contained within our network versus those outside of it, this correlation is higher among morphological variables that are the same or similar to those cell shape features correlated with gene expression modules used to construct the network. The correlation between combined centrality and drug absolute effect on cell area (n=37) was moderate but significant for cytoplasm area, cytoplasm perimeter, nucleus area, nucleus length, nucleus half-width and nucleus perimeter (with Spearman correlation coefficient between 0.34 - 0.37 for all of them, with P<0.05). This correlation in change in morphological features with the centrality of the targeted kinases illustrates the relevance of our constructed network in regulating cell shape. For variables that were not correlated to any gene expression module, we see visibly lower correlation coefficients and insignificant associations (Spearman correlation coefficients of 0.05 - 0.29, P>0.05). These results illustrate that the topology of our network explains some of the variation in the effect of kinase inhibitors tested, in a manner that is feature specific to the ones that were used to construct the network model.

### Network propagation of activated TFs reveals differentially activated processes in the cell shape regulatory network

As transcription factor activity remains the most reliable indicator of signaling that can be extracted from transcriptomics data [51]we applied network propagation to identify sub-networks and nodes of which differentially regulated transcription factors have an effect. The algorithm Random Walk with Restart (RWR [52]) was used to diffuse from activated and inactivated transcription factors in our network reflected by the normalized enrichment scores of transcription factors identified by DOROTHEA [45] (Figure 5A & B; Supplementary Table 9; Methods).

**Figure 5:**
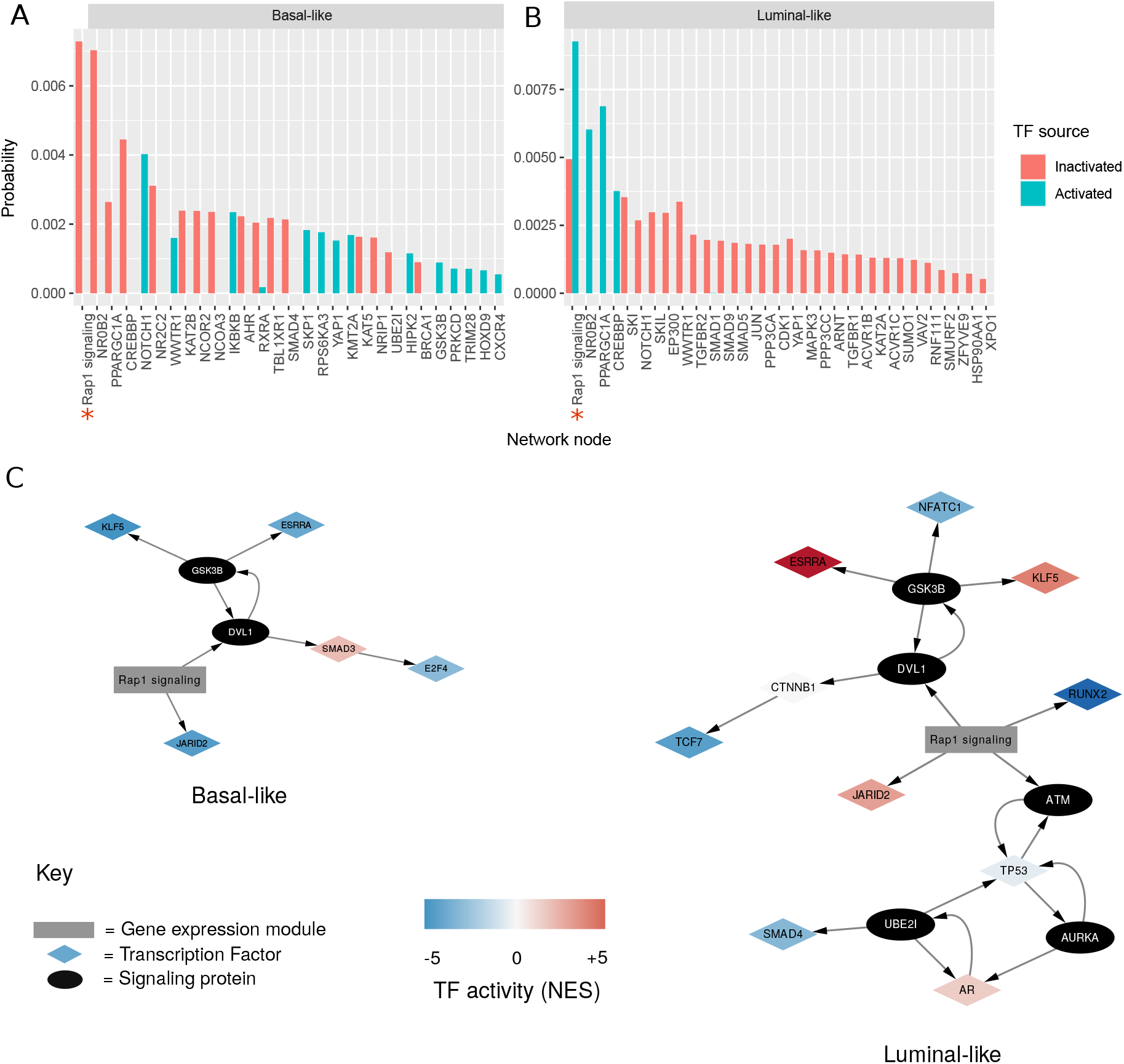
Bar plot showing network propagation in predicted cell shape network from activated and inactivated transcription factors in basal-like cell lines **(A)** and luminal **(B)**. The y axis is a steady state probability (or the ‘heat’ of the nodes in the network after the diffusion) over the graph imposed by the starting seeds, ordered by size. Red bars represent propagation from transcription factor seeds that are predicted to be in-activated, and blue bars show propagation from transcription factor seeds that are predicted to be activated. Red stars along the x axis indicate supernodes that represent gene expression modules. Only those nodes with combined probability > 0.0001 are shown, with the full results available in Supplementary Table 9. **(C.)** Sub-networks illustrating the paths between activated transcription factors (in basal-like and luminal-like) and the ‘RAP1 signaling’ gene expression module. Transcription factors are shown as diamond-shaped nodes, with their colour representing their activity. The ‘RAP1 signaling’ gene expression module is shown as a grey rectangle. Signaling proteins are shown as black nodes.

The most relevant super-node in both luminal and basal diffusions was the gene expression module, RAP1 Signaling, a module which is correlated with several cell shape variables (neighbour frequency, ruffliness, nuclear by cytosolic area and cell width to length) and is enriched in members of the mechanosensitive RAP1 signaling pathway. By performing RWR diffusions on each of the seed nodes separately (Figure S5A-B) we can see that the source of this module’s probability is from the transcription factors JARID2 and RUNX2 in luminal-like cell lines, and JARID2 for basal-like. However, the transcription factors KLF5 and ESRRA in both morphological subtypes also contribute to the ranking of RAP1 signaling, via GSK3B and DVL1 (Figure 5C).

Specific proteins that were top ranked after performing the network propagation in basal-like cell lines include the orphan nuclear receptor NR0B2. Individual RWR found 3 seed transcription factors responsible for this node’s high probability: AR, ESRRA and NR1H3. Other proteins flagged by the propagation were SMAD4, which is regulated by TGF-*β*, IKBKB, which is an activator of NFkB and YAP1. For luminal-like cell lines, NR0B2 is also significantly ranked from the network propagation (as a result of ESRRA activity) as well as transcriptional co-activator PPARGC1A and CREBBP.

### RAP1 gene expression module correlates with known morphologically-relevant TFs in both cell culture and clinical samples

To explore the significance of the RAP1 gene expression module in breast cancer we measured its activity (Methods) in 78 BRCA cell lines. This enabled the correlation of its combined activity with the activity of known transcription factors predicted by DOROTHEA [53] (Figure S6). We find that RAP1 GEM activity was significantly correlated (Kendall; P<0.01, FDR adjusted) with the activity of 19 TFs. Among these are RUNX2 (consistent with the results from our network propagation), TEAD1 (TF mediating the function of YAP1/TAZ) and NFkB. We also correlated transcription factor activity using the same method on tumour samples extracted from TCGA (https://www.cancer.gov/tcga). Using this publicly available dataset, we studied 1,090 BRCA tumours and performed differential expression on each sample. We found 40 TFs significantly correlated (Kendall; P<0.01, FDR adjusted) with RAP1 GEM (Figure 6A-B). The intersection of this analysis between in cell lines and the clinical data were the TFs: SP3, NFKB1, ZNF589, ZC3H8, HIF1A, STAT1, ZNF584, ZNF175 and KLF5.

**Figure 6:**
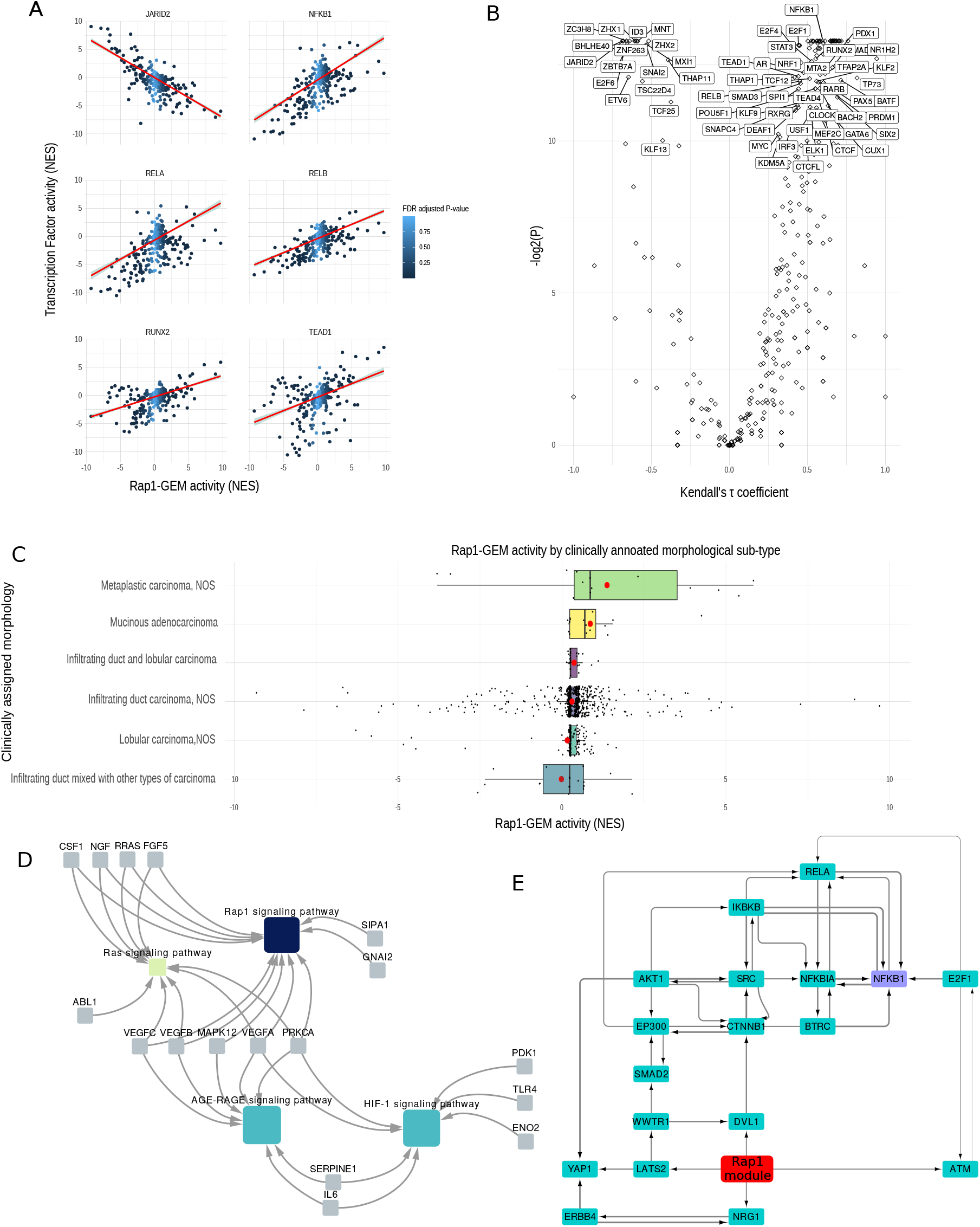
**A** Plots showing the correlation between the RAP1 gene expression module activity (Normalised enrichment score - see methods) and the activity (NES) of various transcription factor (JARID2, NFKB1, RELA, RELB, RUNX2 and TEAD1). Line of best fit according to linear regression is shown in red, with confidence interval in grey. Colour of the points in the plot represents the FDR adjusted P-value of the RAP1 NES as calculated by DOROTHEA. **B**. Volcano plot illustrating the correlation (Kendall’s rank correlation) between activity of RAP1 gene expression module and transcription factor activity, with Kendall’s tau coefficient along the x axis and -log2(FDR adjusted P) along the y axis. **C**. Barplot showing RAP1-GEM activity across different breast cancer samples, separated by clinically assigned morphology. The y axis shows RAP1 gene expression module activity as calculated by DOROTHEA in NES. Mean values for each group are shown by a red dot. **D**. Network showing gene set enrichments of the contents of RAP1 gene expression module. Genes are shown in pale blue, and pathways are shown by nodes whose colour indicates significance of the associated term (-Log2(P)). **E**. Sub-network showing the top flow-carrying edges (99th percentile) calculated using the maximum-flow algorithm between RAP1 gene expression module and NFKB1.

We studied the expression of the module in tumour samples and compared different groups of clinically annotated morphological subtypes. The morphological subtype with the highest overall RAP1-GEM activity was metaplastic carcinoma, a subtype characterised by poorly cohesive sheets [54] and a high propensity to metastasize [55] (Figure 6C). This morphological subtype has a distribution significantly greater (P < 0.005; Two sample Kolmogorov-Smirnov test) than the most frequently assigned morphological subtype (Infiltrating duct carcinoma, NOS). This subtype is a common and homogenous breast cancer grouping characterised by its failure to exhibit morphological features that might allow it to be classified as anything more specific [56].

### Content of RAP1 GEM and its network-neighbourhood allow us to explore potential signaling events relevant in the regulation of cell shape

To understand latent processes driven by components within our gene expression module, we also studied interactions between the gene ontology terms enriched within RAP1 GEM (Figure 6D). This revealed that, as well as RAP1 signaling, the GEM is enriched in AGE-RAGE signaling pathway and HIF1 signaling pathway (consistent with HIF1A’s activity correlating highly with RAP1 GEM in both cell line patient data). HIF1 is known to be regulated by RAP1 [57], although not explicitly in breast cancer.

NFkB has been previously linked to the regulation of cell shape in breast cancer. To explore the interface of RAP1 GEM with NFkB in terms of intra-cellular signaling, we identified a subnetwork of our network responsible for mediating ‘information-flow’ between those two nodes, using the algorithm maximum flow (Figure 6E). By studying the flow of information from RAP1 signaling, we can see that a LATS2/WWTR1/DVL1 (all of WNT signaling) lies between the target and source nodes with much of the flow being carried via these edges. This implicates YAP1/TAZ as being a key effector of the identified gene expression module. This finding is supported by TEAD1 (mediating gene expression of YAP1 and WWTR1/TAZ) being among the most highly correlated of TFs with RAP1 GEM (Figure 6B and S6A).

## Discussion

We present a method that uses transcriptomics and phenotypic data to derive a concise sub-network describing the signaling involved in the regulation of cell shape. This analysis recovered known processes like ‘adherens junction proteins’, ‘cadherin’ [58]and ‘integrin’ [59] [60] as well as pathways responsible for the regulation of cell shape in development, such as WNT [61][62], TGF-*β* [63] and NOTCH [64]. All of these pathways have previously been linked to the development of metastatic phenotypes in breast cancer cells [65] [64] [66]. Importantly, this analysis also sheds light on processes with less characterized associations with cell shape in cancer. We found that a gene expression module enriched in RAP1 signaling, is significantly correlated with cell shape, and is the most differentially expressed module between Luminal-like and Basal-like cell line clusters. We found that it was upregulated in basal-like cell lines while downregulated in luminal, consistent with its negative correlation with neighbour fraction; a cell shape feature most contributing to the ‘cobblestone’ like features of an epithelial and non-metastatic cell type. This gene expression module was also an important node in our identified signaling network, being at the network confluence of multiple activated transcription factors. We also showed this gene expression module to be expressed in patient data, with its activity being correlated with known developmental and morphologically related transcription factors, as well as those used to identify it in the network propagation analysis. In this way,our methodology uses cell line data for network construction and validation, but through our network approach we focus in on more general effects which can be tested and successfully validated in a wider breast cancer clinical context. Hence, we believe these results to be relevant in more general breast cancer applications, but are also sensitive to the inherent context-specificity that exists in biology.

The name-sake of our identified module; RAP1 is a small GTPase in the Ras-related protein family that has been shown to be involved in the regulation of cell adhesion and migration [67][68]. Specifically, RAP1 has been shown to modulate and activate NFkB activity in response to TNF-*α* stimulation in mesenchymal stem cells [69] and modulate migration and adhesion [70]. RAP1 is able to regulate IKKs (IB kinases) in a spatio-temporal manner [71], and is crucial for IkBK to be able to phosphorylate NFkB subunit p65 to make it competent [72] Here, we used our network-centric methodology to highlight a transcriptomic module, characterised in part by RAP1 signaling and that this is a key node in our phenotype-specific signaling network. It is possible that our observations of the significance of RAP1 are as a result of more ‘direct’ interaction between RAP1 and the cytoskeleton. However, the transcriptomic module which we observe accounts for a much larger system-wide rewiring than simply the modulation of cytoskeletal proteins. This implies more complex transcriptional changes that are characteristic of a more robust breast cancer niche.

The RAP1 signaling GEM identified in the network analysis represents a subset of the transcriptome observable among our analysed cell lines. While it is enriched in RAP1 signaling, it is important to note that it represents a collection of latent biological processes rather than a single pathway assigned to it by gene set enrichment. From our network analysis we hypothesise that it is able to interact with intracellular signaling pathways in order to modulate transcription factor activity and consequently cell shape. Other pathways enriched in the expression module include HIF-1 signaling pathway, which is known to be activated by RAP1 in melanoma [57], but this has not been shown in breast cancer. Also, AGE-RAGE signaling was also enriched in our module of interest. AGE-RAGE signaling pathway has recently been shown to overlap with RAP1 signaling pathway in cardiac fibroblasts to alter the expression of NFkB [73], although this crosstalk has also not been illustrated in breast cancer. Here, we observe genes of all these pathways functioning as a cohesive unit in breast cancer in a previously unobserved fashion. Additionally, their combined expression correlates with morphological traits associated with a negative prognosis.

We also observe that our gene expression module of interest is significantly correlated with NFkB in both clinical samples and cell culture. Other authors have flagged the direct effect of RAP1 on the cytoskeleton and NFkB [74][69], but here we go further, using our unbiased systems approach to link RAP1 signaling with multiple transcription factors and pathways. Based on known functions of RAP1, along with the functions of pathways that we find interact with it, we hypothesize that the identified transcriptomic unit is key in relaying information from a cell’s physical environment to modulate and maintain the cancer stem cell niche [75].

Previous studies have established a connection between the NFkB signaling pathway and regulation of cell shape in breast cancer [26][25]. Our findings also illustrate the significance of this pathway in the regulation of cell morphology, with multiple NFkB regulators and transcriptional co-activators being flagged in our results. Some morphology-correlated gene expression modules were significantly differentially expressed between cell shape subtypes with the ARNT KO module being significantly upregulated in basal-like cell shapes relative to luminal. We also found this gene expression module to have the highest total correlation with all of the morphological features, indicating a strong association with cell shape. By studying terms enriched in this module from the EnrichR library, we find both ‘TNF-*α* signaling via NFkB’ to be enriched as well as genes downregulated during AHR nuclear translocator (ARNT) shRNA KO. Signaling by TNF is able to activate NFkB, a transcription factor known to control the expression of many EMT related genes [76] which has shown to be more sensitive to TNF-a stimulation in mesenchymal-like cellular morphologies than epithelial. This was hypothesised to generate a negative feedback which reinforces a metastatic phenotype of breast cancer cells [25]. Here we observe also that an ARNT KO/TNF module is upregulated in basal-like cell lines, consistently with these findings. ARNT is a protein shown to be involved in regulating tumour growth and angiogenesis along with its binding partner aryl hydrocarbon receptor (AHR) [77]. Previous studies have also shown its ability to modulate NFkB signaling with the activated form possibly interfering with the action of activated p65 [78]. Our findings that the upregulation of a gene expression module that is associated with ARNT knockdown further gives credence to NFkB being positively regulated in mesenchymal-like cell morphologies. Furthermore, the results of our network propagation yielded activators and transcriptional coactivators of NFkB (IKBKB [72], NR0B2 [79] and CREBBP [80]. These findings indicate that NFkB is modulated by both phosphorylation (through stimulation by TNFa), spatial-temporal location (through RAP1) and transcriptional co-activation (through NR0B2 and CREBBP) in breast cancer in a shape-dependent manner.

Aside from the biological findings of this study, we illustrate an approach for network analysis of a specific course-grained phenotype through expression; a notoriously poor (if cheap and widely available) proxy for gauging intracellular signaling [81]. In contrast to existing methods that use gene expression as a direct proxy for signaling [82][83][84][85], our approach infers transcription factor activities from the expression data and uses these as an anchor to infer upstream signaling networks relevant to the regulation of our phenotypes. Transcription factor activities can represent the outcome of a signal transduction process compared to the expression profiles and are thus a better proxy for cell signaling activities of the cell [86]. Such an approach has been previously used, for example by the tool CARNIVAL [87]. However, this and other available tools neglect the propensity for the transcriptome to be regulated in a highly context specific and modular structure [88][89]. Moreover, their reliance on annotated pathways to describe cell signalling undermines their ability to spot novel functional units, specific to a given phenotype. Here, using context-specific gene expression modules, we produced a network connecting the genes of interest from diverse analyses and used a network propagation algorithm to further focus on signaling proteins of novel interest. While there inevitably remains a level of bias stemming from the transcription factor regulon and pathway annotations, our bottom-up approach seeks to identify latent modular structure within transcriptomic data first. This puts the emphasis on data-driven gene expression modules, rather than literature-derived regulons and pathways. This approach takes an important step towards reducing the bias associated with previously annotated pathways and allows the identification of important regulatory units and their function with respect to cell shape from a systems biology point of view. Our network approach allows us to map the interface between two graphically presented systems in the cell; the transcriptome and intracellular signaling. Both can be easily combined with complex, multivariate phenotypic data which here has revealed a clearer picture of how signaling regulates cell morphology in breast cancer.

The interoperability of this approach is obvious, with any number of continuous variables measured with gene expression able to be correlated with module eigengenes using WGCNA. Here, we used OmniPath as a base network, but other network-based representations of the cellular environment can be used based on the appropriate context. Thus, our method represents a data-driven, network-based approach compatible with many different multi-scale phenotypes that are driven by intracellular signaling.

Overall, our unbiased network-based method highlights potential ‘missing links’ between sensing extracellular cues and transcriptional programmes that help maintain the cancer stem cell niche, and ultimately push breast cancer cells into EMT and metastasis. These represent starting points for further experimental studies to understand and therapeutically target the links between cell shape, cell signaling and gene regulation in the context of breast cancer.

## Materials and methods

### WGCNA analysis

Using Weighted correlation network analysis, we performed co-expression module identification using the R package WGCNA [33]. We used bulk RNA-seq data from Expression Atlas (in FPKM - E-MTAB-2770 and E-MTAB-2706) acquired from commonly used cancer cell lines of various cancer types [90]. We collated 13 breast cancer and 1 non-tumorigenic cell line for which imaging data was available (BT474, CAMA1, T47D, ZR75.1, SKBR3, MCF7, HCC1143, HCC1954, HCC70, hs578T, JIMT1, MCF10A, MDAMB157 and MDAMB231 [25]. We acquired representative images of each cell line from Sero et al., 2015 (https://datadryad.org/stash/dataset/doi:10.5061/dryad.tc5g4). Cell imaging segmentation was performed using Acapella software (PerkinElmer) with an automated spinning disk confocal microscope. The presented images (Figure 2) are taken from the above link, stained with DAPI (blue), Alexa 488 (green) and DHE (red). Using Ensembl-Biomart, we filtered genes to only include protein-coding genes [91] and genes whose FPKM was greater than 1, leaving a total of 15,304 genes.

We created a signed, weighted adjacency matrix using log2 transformed gene expression values and a soft threshold power (*α*) of 9. We translated this adjacency matrix (defined by Eq.1) into a topological overlap matrix (TOM; a measure of similarity) and the corresponding dissimilarity matrix (TOM - 1) was used to identify modules of correlated gene expression (minimum module size of 30).

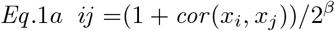

We took morphological variables referring to breast cancer cell lines from Sero et al., 2015, which include 10 significant features shown to be predictive of TF activation [25]. We correlated these features with module eigengenes using Pearson correlation and we tested these values for significance by calculating Student asymptotic p-values for given correlations. Multiple hypothesis testing was performed using a permutation based procedure whereby we recalculated the correlation matrix 1,000 times with resampled data. We then generated null distributions for each ranked correlation statistic in our matrix, and compared them to our real data of the same rank. We include in the Supplementary Table 1 confidence intervals of our permutation-based multiple-correction procedure. For the modules that correlated with morphological features (Pearson Correlation Coefficient 0.5 and Student P<0.05), we identified enriched signaling pathways using the R package EnrichR [34], and the signaling database Reactome [43]. Using the database TRRUST v2 (Accessed : 01/07/18) [35], we identified TF regulons that significantly overlap (Exact Fisher’s test, P<0.1) with the gene expression module contents. This was done separately for inhibitory and activatory expression regulons for each transcription factor, with regulatory relationships of unknown sign being used in the significance calculations for both.

We named gene expression clusters using significantly enriched terms identified by the EnrichR analysis (Supplementary Table 3). As some clusters were very obscure, we utilized the entire EnrichR list of libraries (https://maayanlab.cloud/Enrichr/#stats for full list) with precedence going to the signaling databases of KEGG, Reactome, Panther and Wikipathways (Accessed : 01/04/20) [92][93][48]. Some modules could not be assigned informative terms and so were named ‘not annotated’ (NA).

### Clustering and differential expression

Using the k-means algorithm, we classified the 14 breast cancer cell lines by the median values of each of their shape features (k=3, see Figure S2A). We performed differential expression analysis using the R package DESeq2 [94]. We filtered genes so that only protein coding genes and those with more than 0.5 counts per million in at least 8 cell lines were included. We calculated Log2 fold changes with the cluster of interest as the numerator and the remaining cell lines acting as a control. Using the R package FGSEA [95], we performed gene set enrichment analysis of the differentially regulated proteins using the complete pathways gene set (Release 01 April 2020) from MSigDB [96] and the WGCNA gene expression modules identified in previous analysis. We calculated transcription factor regulon enrichment using the software DOROTHEA (Accessed : 01/04/20) [53].

### Network Generation

Using a Prize Collecting Steiner Forest (PCSF) algorithm, we generated a cell-shape regulatory network implemented through the R package PCSF [46]. For the prize-carrying nodes to be collected by the PCSF algorithm, we used the transcription factors significantly regulating the WGCNA modules using TRRUST (p<0.1), the differentially activated transcription factors identified by DOROTHEA (p<0.1), and the signaling proteins included in the REACTOME pathways that were enriched in transcription factors identified (p<0.05). We identified these pathways by using the TRRUST TFs identified in the previous steps, as well as ENCODE and ChEA Consensus TFs from ChIP-X [39], DNA binding preferences from JASPAR [40][41],

TF protein-protein interactions and TFs from ENCODE ChiP-Seq [42]. Using EnrichR, we identified pathways that were enriched in the identified TFs, and the proteins that were included in these pathways were extracted from Pathway Commons using the R package paxtoolsr [97]. This was tested for bias to specific pathways by generating pathway-specific null distributions from 1,000 resampled GEMs. Distributions of p values for each Reactome pathway were generated, where failed tests (because of no TF enrichment) were given a p-value of 1. Results of this were corrected for multiple-hypothesis testing using FDR correction.

The ‘costs’ associated with each edge in the regulatory network were the inverse of the number of sources linked to each regulatory connection scaled between 1 and 0, such that the more the number of citations for an edge, the lower the cost. For the base network used by the algorithm, we used the comprehensive biological prior knowledge database, Omnipath (Accessed : 06/05/20) [98], extracted using the R package OmnipathR [47]. We set each prize for significant TFs or signaling pathways to 100 and used a random variant of the PCSF algorithm with the result being the union of subnetworks obtained on each run (30 iterations) after adding random noise to the edge costs each time (5%). The algorithm also includes a hub-penalisation parameter which we set to 0.005. Other parameters include the tuning of node prices (set to 1) and the tuning of trees in the PCSF output (40).

We included the WGCNA modules themselves as super-nodes in the network, by adding incoming edges from the transcription factors contained within the regulatory network whose regulomes (as described in TRRUST v2 [35]) significantly overlap (Fisher’s exact test; P<0.1) with the gene content of the module in question. We represented the respective cell-shape phenotypes as nodes in a similar fashion, by including undirected edges from expression modules and phenotypes where there was significant correlation (|PCC| > 0.5 & P<0.05) between them. To account for expression modules’ effect on upstream signaling, we added edges from the WGCNA modules back up to proteins that were themselves included within the modules. We set the edge weight of these to 1, such that any predicted activity of the gene expression module would be translated directly into its constituent signaling proteins and thus account for feedback between cell shape signaling networks, and the context-specific expression modules identified in the first step. We identified enriched terms in the network using the 2016 release of the database Panther [48] and GSE package EnrichR [34].

### Network propagation of functional TFs

We examined the potential effect of significantly activated (FDR < 0.05), and deactivated TFs in different cell line clusters using network propagation in our generated network. We replaced edge weight with Resnik Best Match Average (BMA) semantic similarity [99] between the biological process GO terms of the two interacting pairs, with the sign of the interaction being inherited from Omnipath [47]. We then scaled the semantic similarity edge weights between 1 and -1.

We used the differentially activated transcription factors identified using DOROTHEA (P<0.05) as seeds for a Random Walk with Restart (RWR) algorithm using the R package diffuseR (available at: https://github.com/dirmeier/diffusr). We judged a node to be significantly ranked if its affinity score relative to the inputted seeds was greater than the same node’s affinity score with a random walk simulation performed with randomised seeds. We performed this randomised simulation 10,000 times, from which a p-value was determined to judge significance (P<0.1). We performed this propagation by RWR for both luminal-like and basal-like morphological clusters on significantly activated and deactivated transcription factors separately, in addition to simulations where each seed was considered in isolation. We generated a graphical representation of the network edges and TFs responsible for the ranking of RAP1 signaling by plotting all the shortest paths between RAP1 and the TFs that caused it to have a non-zero affinity score when each TF was considered in isolation.

### Breast cancer cell morphology following kinase inhibitor treatment

We used small molecule kinase inhibition data from Harvard Medical School (HMS) Library of Integrated Network-based Cellular Signatures (LINCS) Center [100], which is funded by NIH grants U54 HG006097 and U54 HL127365 (available from: https://lincs.hms.harvard.edu/mills-unpubl-2015/, Accessed : 01/08/20). This dataset is derived from the treatment of 6 cell lines with a panel of 105 small molecule kinase inhibitors. They measured textural and morphological variables following treatment by high through-put image analysis [49][101]. We combined this assay with another dataset from HMS-LINCS; a Target Affinity Spectrum (TAS) for compounds in the HMS-LINCS small molecule library measuring the binding assertions based on dose-response affinity constants for particular kinase inhibitors (https://lincs.hms.harvard.edu/db/datasets/20000/, Accessed : 01/08/20). Using this dataset, we filtered for only molecule-binding target pairs with a binding class of 1 (representing a Kd <100nM affinity). Further to this, we removed molecules which had more than 5 targets with a Kd of 100nM. Consequently, the remaining kinase inhibitors were relatively narrow spectrum, thus simplifying analysis of their phenotypic effect. We expressed these results as batch-specific log fold changes of 10um drug treatment relative to the mean of the control set (untreated and DMSO treated cells). Spearman rank correlation was calculated between the drug target’s network centrality and the absolute log fold change of the morphological variable. We also used the Kolmogorov-Smirnov statistic to assess significance between cell morphology after treatment with drugs targeting kinases inside versus outside our predicted network. This was also repeated on other breast cancer cell lines and using a TRAIL (apoptosis inducing) control (Supplementary table 8).

The morphological data in the kinase inhibition screen was measured using two dyes (DRAQ5 and TMRE), the intensity of which we used to normalise textural features and the measurement of cytoplasmic and nuclear small spots. We reported counts for small nuclear or cytoplasmic spots as a mean of the individually normalised readings from both dyes. We considered a treatment perturbing our network if at least one of the kinase inhibitors targeted a protein that was represented by a node within the network.

### Quantifying kinase inhibitor influence

We incorporate information from the Target Affinity Spectrum assay, as well as graph-based properties of kinase inhibitor targets, using the product of the Szymkiewicz-Simpson similarity (measured between the cell shape network nodes and the drug targets) and the centrality of the targeted nodes in the predicted network with semantic similarity edge weights. The product of these generates, for a given kinase inhibitor the statistic:

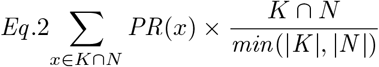

Where K is the set of kinases an inhibitor is predicted to target, N is the nodes of the network and the function PR() is the centrality of a particular node in the network as defined by the PageRank algorithm [102]. This centrality measure has been shown to be effective in prioritizing proteins by relative importance in signaling or protein-protein interaction networks [103]. We used this statistic as a measure of a kinase inhibitor’s influence on cell shape.

### Analysis of BRCA cell line and TCGA sample RNAseq data

For the cell lines, we used RNA-seq data from Expression Atlas (in FPKM - E-MTAB-2770 and E-MTAB-2706) [90]. This was analysed using DESeq2 [94] as per the methodology in the subsection entitled “Clustering and differential expression”. Both TF and module activity was calculated using the algorithm VIPER [104]. For patient data, the results shown here are based upon data generated by the The Cancer Genome Atlas (TCGA) Research Network:https://www.cancer.gov/tcga, Accessed : 01/04/21. For computational efficiency, we use Gamma-Poisson models to predict differentially expressed genes from our samples using the package glmGamPoi [105]. We use the sample of interest as the numerator with the remaining tumour samples acting as a control. For quantifying correlation between RAP1 - GEM and different transcription factors we remove samples with insignificant activation of either the TF in question or RAP1-GEM (FDR adjusted P value < 0.05). Correlation was quantified using Kendall rank correlation coefficient. Differences in distributions of morphological subtypes was quantified by Kolmogorov-Smirnov test.

### Maximum-flow network analysis

For maximum-flow calculations, we used the Resnik BMA semantic similarity [99] as the maximum ‘carrying capacity’ of an edge in the network. To visualise the optimised solution (as implemented by the R package igraph [106] we selected only those edges in the 99th percentile of the flow-carrying edges in the network. Visualisation was performed using the software, Cytoscape [107]. Maximum flow was performed with the R package igraph [106].

### Quantification and Statistical Analysis

Statistical tests were performed in base R unless otherwise mentioned in the methods and p-value cut-offs are shown in parentheses after reporting an effect as significant. Weighted Pearson correlation with t-test for significance was used to correlate eigengenes and cell shape features using the RNA package WGCNA. We used a one-way ANOVA test for comparing the means of the shape variables among the identified 3 cell line clusters (n = 75,653) and a Tukey honest significant differences test to perform multiple-pairwise comparison among the means of the groups. The same tests were performed on the differences in 10 cell shape variables when HS578T was treated with 37 kinase inhibitors (n = 23,128). Fisher exact test was used to test significance of overlap between TRRUST regulons and identified gene expression modules (Supplementary Table 4 shows the size of the overlap).

Enrichment of gene sets was performed by EnrichR, an enrichment library that utilises a hyper-geometric test to identify significantly enriched terms in a gene list. This tool (described in [108] calculates a score combining the Fisher exact test p-value of the enrichment with the z-score deviation from the expected rank. The pre-ranked gene-set enrichment algorithm FGSEA was used for the identification of enriched terms in the differentially expressed genes allowing for accurate estimation of arbitrarily low P-values that occur in expression datasets.

Spearman rank correlation was used to measure the strength of the association between target network centrality and the measured effect of its perturbation by inhibition. Spearman was chosen because the centrality (combined with Szymkiewicz-Simpson) according to equation 2 does not follow an exact normal distribution. Kendall rank correlation coefficient was used when calculating the correlation between TF activity and RAP1-GEM activity because confidence intervals for Spearman’s r_S_ are less reliable and less interpretable than confidence intervals for Kendall’s -parameters. When trying to distinguish between many correlations of similar quality, this becomes more important. FDR adjustment for multiple testing correction was always used when multiple tests were performed in the same analysis.

Kolmogorov-Smirnov test was used to measure differences in distributions of clinically assigned tumour morphologies. This was because clinical groupings are mixed (i.e. Infiltrating duct and lobular carcinoma) and others are characterized by an absence of features over their presence. This means that the assumption of normality required for a t-test is not fulfilled.

For differential expression analysis the DESeq2 R package [94] was used. DESeq2 fits negative binomial generalized linear models for each gene and uses the Wald test for significance testing. The package then automatically detects count outliers using Cooks’s distance and removes these genes from analysis.

Significance was determined for RWR network propagation by randomising seed nodes (preserving their values) 10,000 times and selecting only the non-seed nodes that were significantly ranked relative to the randomised simulations (P<0.1).

## Data and Software Availability

The complete R scripts used for this methodology are available on Gitlab; https://gitlab.ebi.ac.uk/petsalakilab/phenotype_networks.

## Acknowledgements

We would like to thank EMBL-EBI for funding the project. We would also like to thank Bishoy Wadie and Vivian Robin for critical reading of the manuscript.

## Author Contributions

**CB:** Conceptualization, Methodology, Software, Validation, Formal analysis, Writing - Original Draft. **EiP:** Conceptualization, Methodology, Software. **GG:** Conceptualization, Supervision. **JS:** Resources. **ENE:** Validation, Writing - Original Draft. **ChrB:** Writing - Review & Editing, Resources. **EP:** Writing - Review & Editing, Conceptualization, Supervision, Methodology, Project administration, Funding acquisition.

