## Supplementary figures for "Identification of phenotype-specific networks from paired gene expression-cell shape imaging data"

A

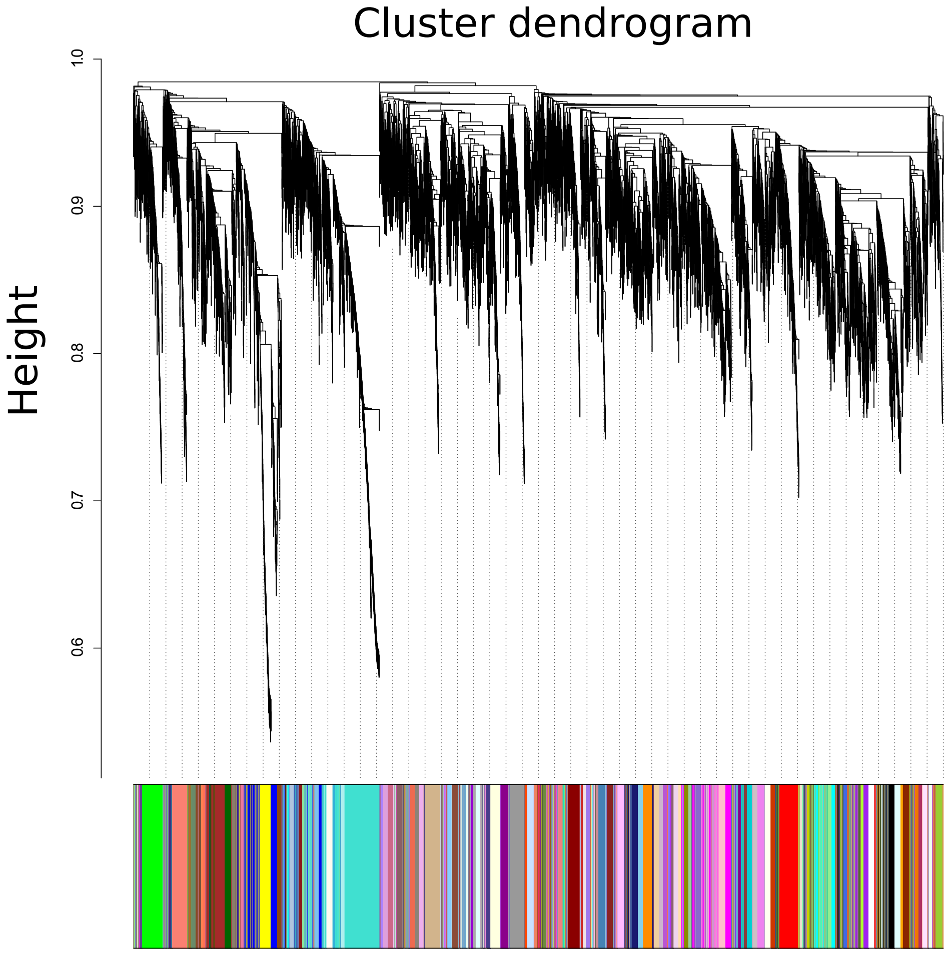


B

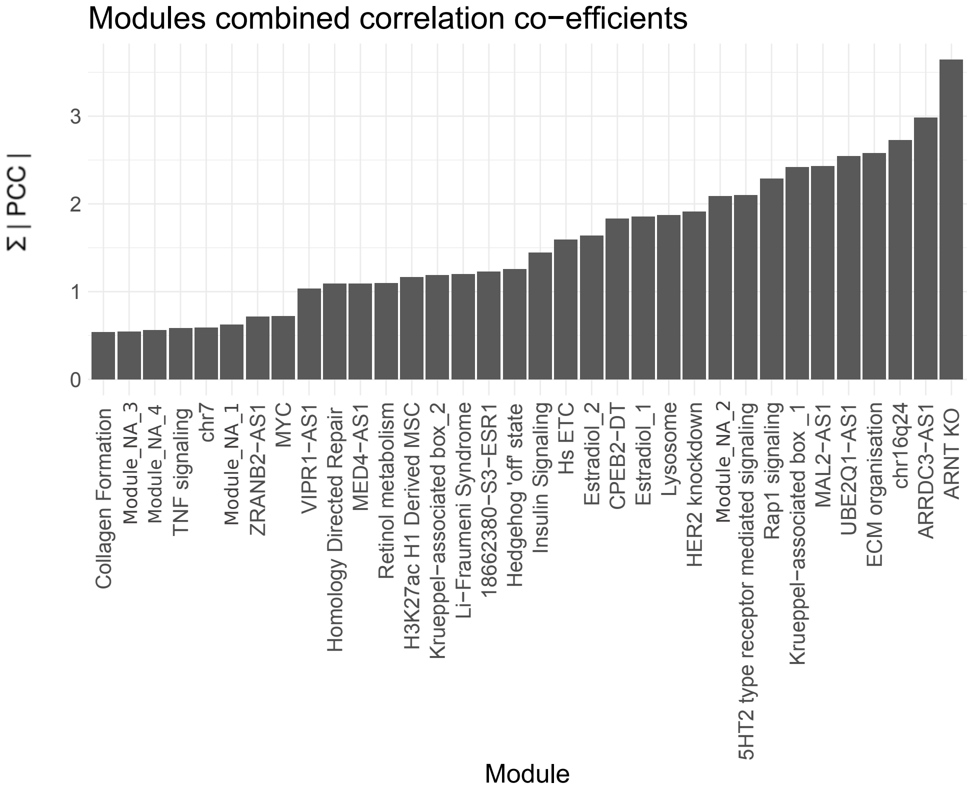


**S1A** - Gene expression dendrogram showing the hierarchical arrangement of gene expression in breast cancer cell lines. Derived gene expression modules are shown along the bottom. **B** - Bar chart showing the sum of the absolute values of Pearson's correlation coefficient (PCC) for the significantly correlated (P<0.05) module - feature associations.

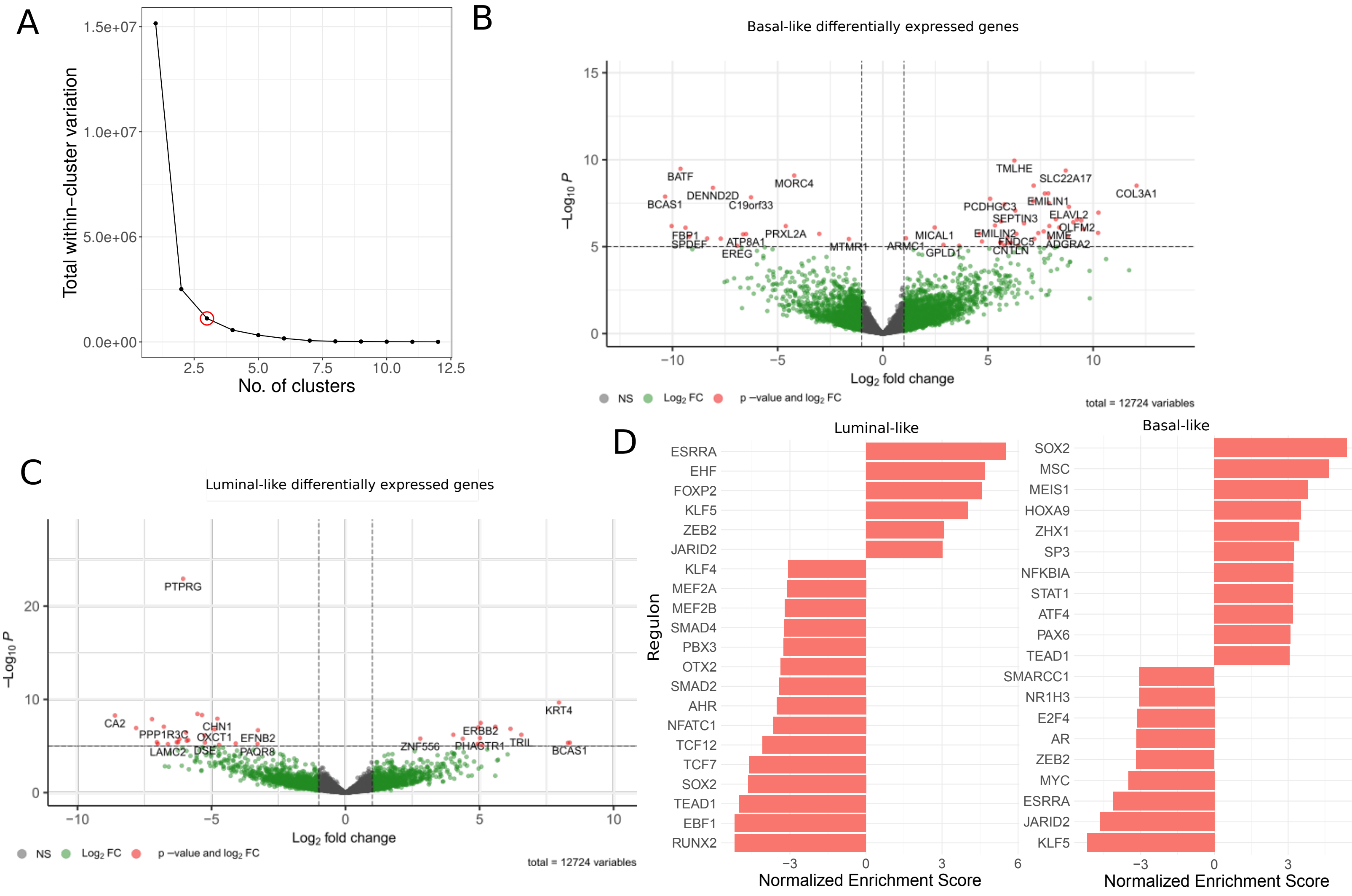


**S2A** - Elbow plot illustrating the diminishing return of decreasing total within-cluster variation of cell shape groups as the number of clusters is increased. A red circle shows the selected statistic for k (3). **B & C.** Volcano plots showing significantly differentially expressed genes for basal-like and luminal-like morphological clusters. **D.** TF activity for morphological clusters C and B (luminal-like and basal-like respectively) as derived from RNAseq.

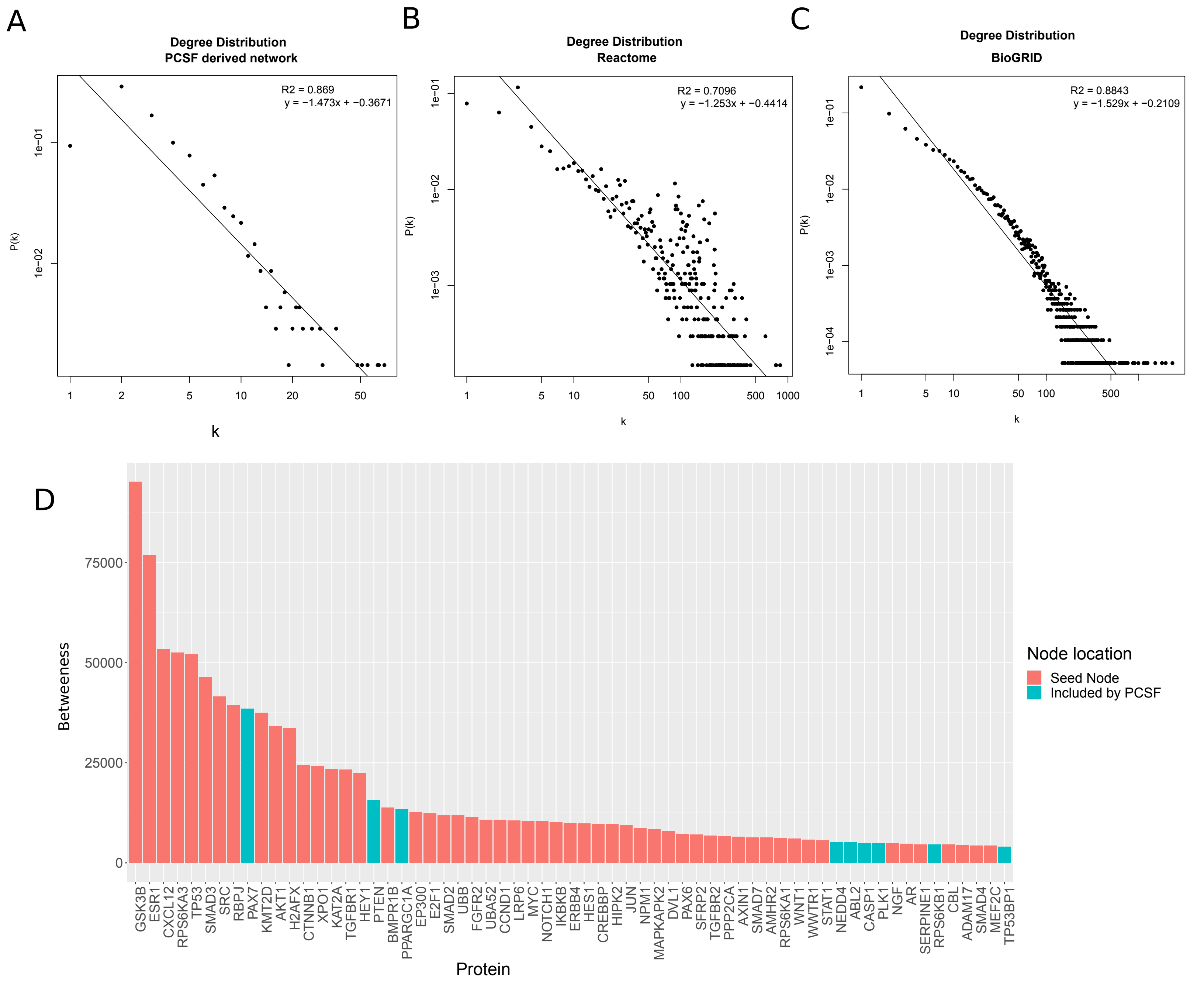


**S3A-C** - Plots showing the scale degree free distribution of our PCSF derived network **(A)**, consistent with the same plots from Reactome **(B)** and BioGRID **(C)** on a log-log scale. The x-axis shows the number of connections for a given vertices (degree - shown by k), while the y axis represents the proportion of vertices in the network having a degree of k. The degree exponent is shown as the gradient of the line of best fit. **D -** Bar chart showing the betweenness centrality of nodes in the derived regulatory network, with centrality on the y axis and the nodes (representing proteins) on the x axis. Nodes are coloured red if they were original seed nodes used in the PCSF and blue if they were included by the algorithm. Only those nodes with centrality > 4,000 are shown.

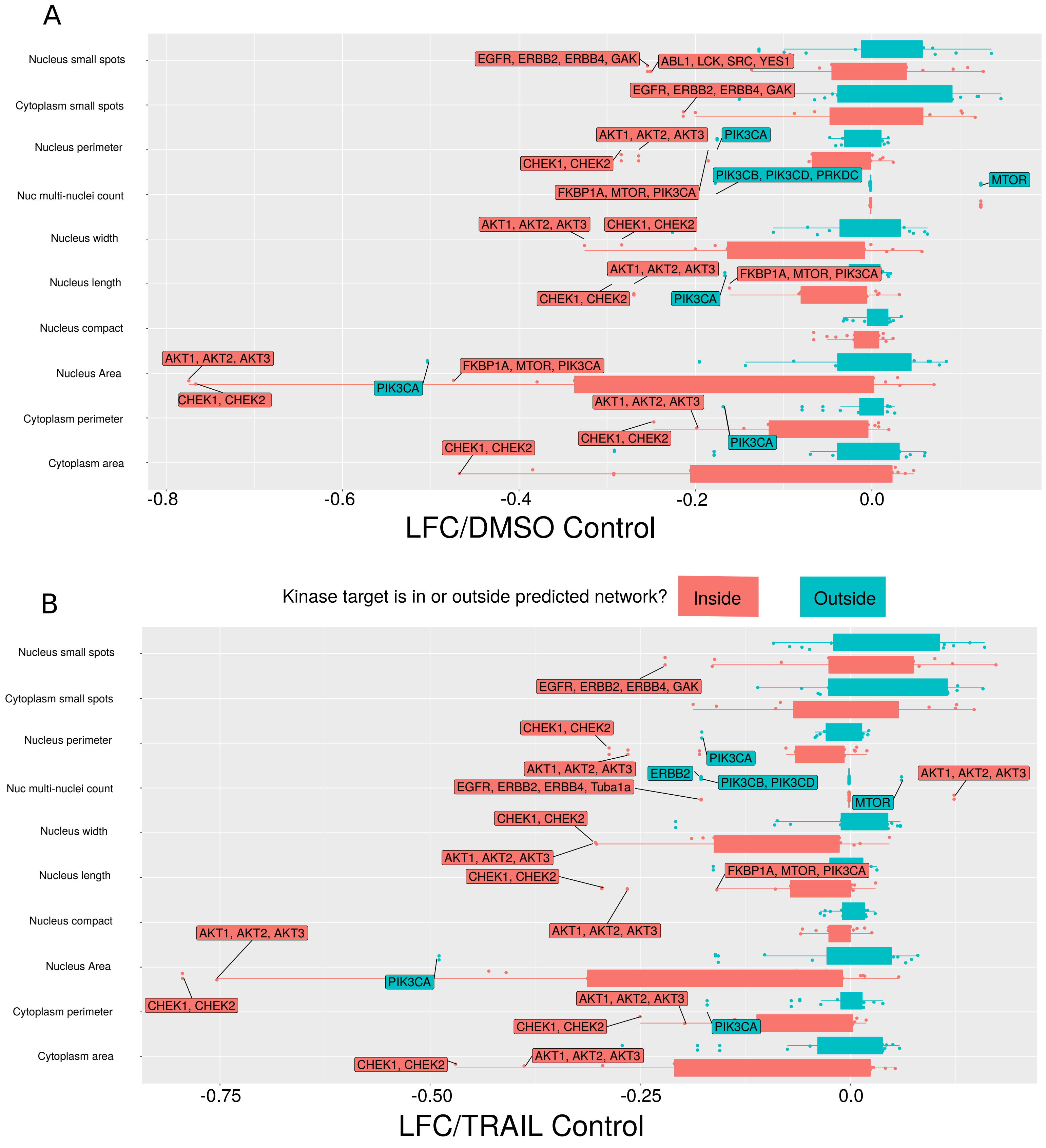


**S4** - Box plots showing the log fold changes of cell shape variables after treatment with a drug relative to a DMSO control **(A)** and TRAIL control **(B).** The kinase targets of the drugs are shown as labels for the outliers (IQR method), with inhibitors targeting drugs within the predicted network coloured red and those targeting kinases not included in our network coloured blue.

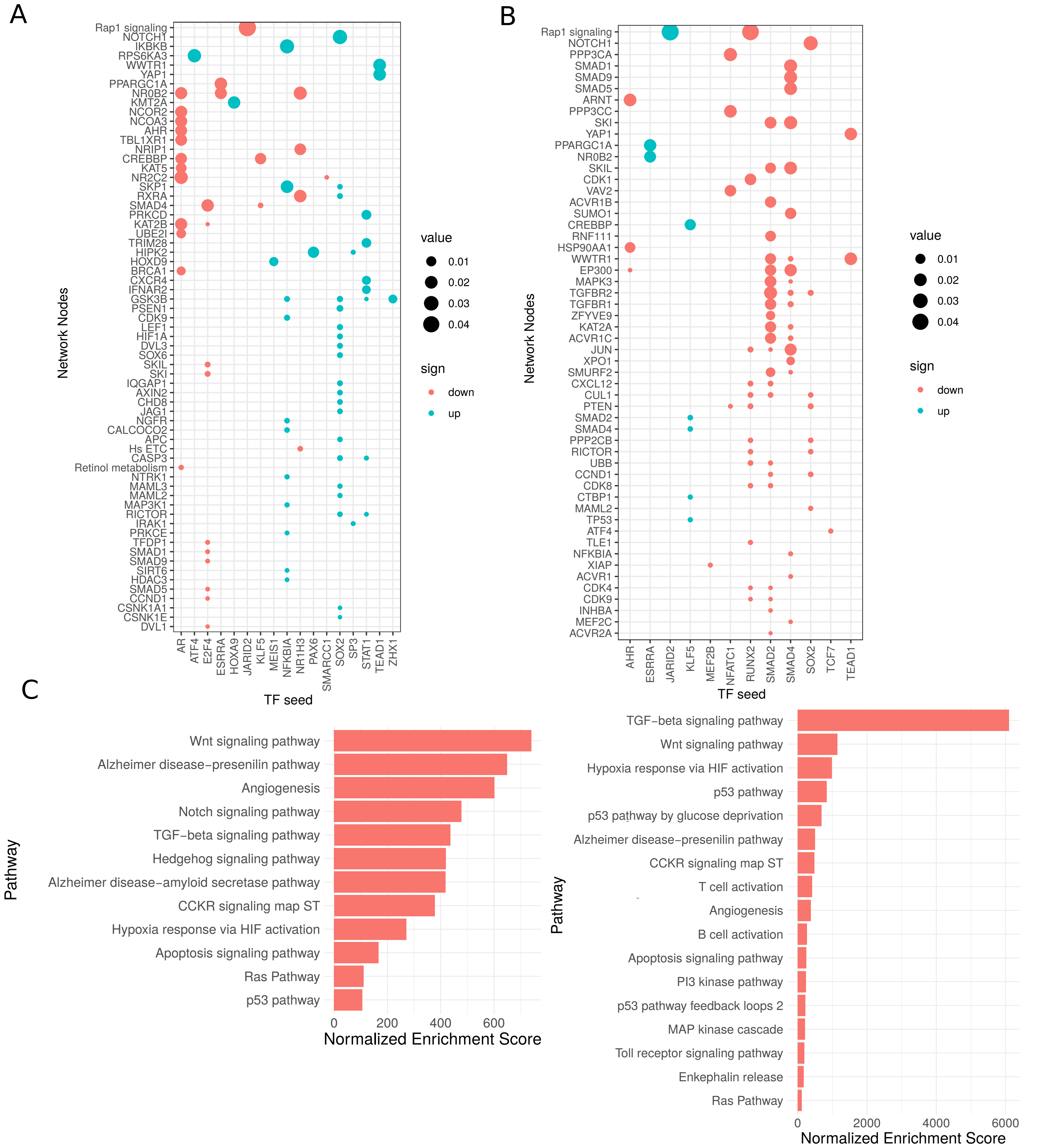


**S5A** - Dot plot showing network propagation in predicted cell shape networks from activated and inactivated transcription factors in basal-likecell lines, performed one seed node at a time. The y axis shows different nodes in the network, with red stars indicating super-nodes(gene expression modules). The x axis shows the TFs used as seed nodes. The size of the dot indicates the steady state probability over the graph imposed by the starting seeds and the colour shows whether or not the TF source was predicted to be activated or deactivated from the gene expression data. **B** - Dot Plot illustrating the network propagation for luminal-like cell lines. **C** - Barchart showing normalised enrichment score (NES) of significantly (adjusted P<0.01, Benjamini-Hochberg) enriched terms associated with high ranked nodes after basal-like TF network propagation. **D** - Bar chart showing normalised enrichment score (NES) of significantly (adjusted P<0.01, Benjamini-Hochberg) enriched terms associated with high ranked nodes after luminal TF network propagation.

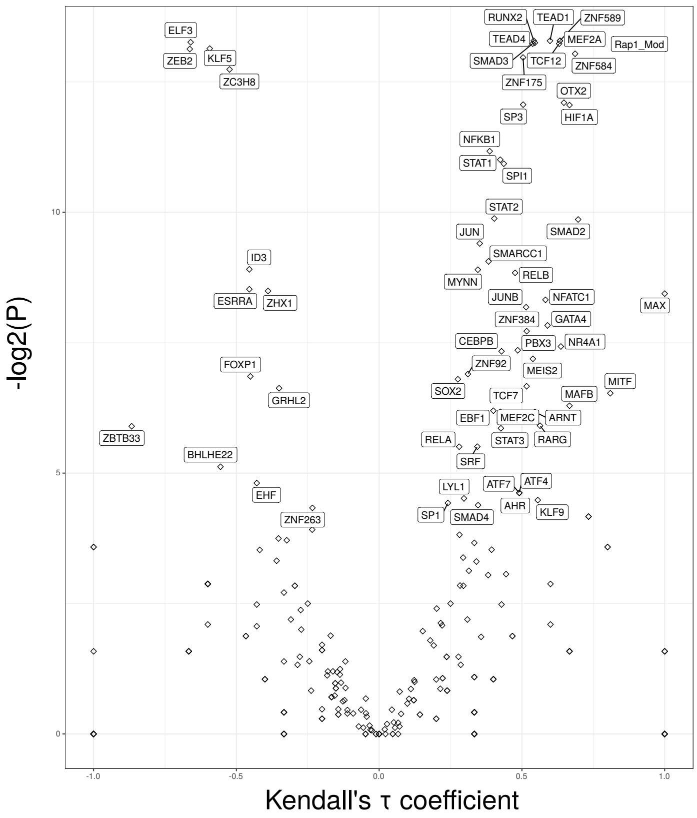


**S6 -** Volcano plot illustrating the correlation (Kendall’s rank correlation) between activity of Rap1 gene expression module and transcription factor activity in 78 breast cancer cell lines. Kendall’s tau coefficient along the x axis and -log2(FDR adjusted P) along the y axis.
